# Disruption of large-scale electrophysiological networks in stroke patients with visuospatial neglect

**DOI:** 10.1101/602318

**Authors:** Tomas Ros, Abele Michela, Anaïs Meyer, Anne Bellman, Philippe Vuadens, Victorine Zermatten, Arnaud Saj, Patrik Vuilleumier

## Abstract

Stroke frequently produces attentional dysfunctions including symptoms of hemispatial neglect, which is characterized by a breakdown of awareness for the contralesional hemispace. Recent studies with functional MRI (fMRI) suggest that hemineglect patients display abnormal *intra-*and *inter-hemispheric* functional connectivity. However, since stroke is a vascular disorder and fMRI signals remain sensitive to non-neuronal (i.e. vascular) coupling, more direct demonstrations of neural network dysfunction in hemispatial neglect are warranted. Here, we utilize electroencephalogram (EEG) source imaging to uncover differences in resting-state network organization between patients with right-hemispheric stroke (N = 15) and age-matched, healthy controls (N = 27), and determine the relationship between hemineglect symptoms and brain network organization. We estimated *intra-*and *inter-*regional differences in cortical communication, by calculating the spectral power and amplitude envelope correlations (AEC) of narrow-band EEG oscillations. We firstly observed focal frequency-slowing within the right posterior cortical regions, reflected in relative delta/theta power increases and alpha/beta/gamma decreases. Secondly, nodes within the right temporal and parietal cortex consistently displayed anomalous intra- and inter- hemispheric coupling, stronger in delta and gamma bands, and weaker in theta, alpha, and beta bands. Finally, a significant association was observed between the severity of left-hemispace search deficits (e.g. cancellation test omissions) and reduced functional connectivity within the alpha and beta bands. In sum, our novel results validate the hypothesis of large-scale cortical network disruption following stroke, and reinforce the proposal that abnormal brain oscillations may be intimately involved in the pathophysiology of visuospatial neglect.

## Introduction

Apart from motor or sensory impairments, the sequelae of ischaemic stroke may cause a significant impact on attentional function. This is most apparent when stroke damage leads to symptoms of hemispatial neglect (Domínguez-Borràs et al. 2013; Vuilleumier and Rafal 2000), which is characterized by an inability to attend to and process information from the left (or more rarely right) side of space (i.e., contralateral to the lesion site). Hence, right-hemisphere stroke patients exhibit an impairment in detecting visual (or auditory) stimuli in their left hemifield (and vice-versa for patients with left hemisphere stroke). Despite the fact that this is a major source of disability in patients’ daily life, current treatments for hemineglect remain minimally effective. Moreover, untreated hemineglect leads to poorer recovery prognosis and reduced benefits from rehabilitation therapies for other neurological deficits.

Therefore, a deeper understanding of this clinical phenomenon is required for several inter-dependent reasons: a) to better identify the neurobiological targets for rehabilitation, b) to determine the underlying physiopathological anomalies, in particular whether local and/or distributed brain dysfunction is crucially implicated, and c) to elucidate the neural mechanisms that give rise to perceptual consciousness.

To address these issues, in line with recent work, here we investigate how brain activity dynamics is altered following focal hemispheric stroke, at both local and global levels, and what is the functional impact of such changes on attentional performance in patients. To this aim, we use EEG recording and adopt a network framework by reducing the whole-brain to a manageable number of regions-of-interest (nodes), whose functional activity is considered to covary with each other through pairs-wise connections (edges). Utilizing this approach, emerging work has suggested that hemineglect might be associated with significant disruptions of brain functional connectivity. Specifically, several functional neuroimaging studies reported that the visuospatial deficits in hemineglect are accompanied with abnormal interhemispheric and/or intrahemispheric connectivity (Baldassarre et al. 2014; Fellrath et al. 2016; Guggisberg et al. 2014; Sasaki et al. 2013; Yordanova et al. 2016). Most recently, using functional MRI (fMRI), Ramsey and colleagues showed that recovery from hemineglect is linked to the return of previously depressed interhemispheric connectivity between nodes of sensorimotor and attention networks (Ramsey et al. 2016). However, since stroke is a vascular disorder and fMRI signals remain sensitive to non-neuronal (i.e. vascular) coupling, stronger validation of neural network dysfunction in hemineglect is warranted. Physiologically speaking, fMRI measures local changes in brain metabolism which generally echo (but lag by a few seconds) electrophysiological activation/deactivation patterns of neuronal populations (Bentley et al. 2016; Hermes, Nguyen, and Winawer 2017; Hutchison et al. 2015). Interestingly, studies comparing non-invasive electrophysiological methods such as magnetoencephalography/electroencephalography (M/EEG) with fMRI have found a tight spatio-temporal correspondence between the amplitude envelopes of neural oscillations and neurovascular fMRI signals (Brookes et al. 2011; Mantini et al. 2007; de Pasquale et al. 2010), although there are several instances of dissociations between fMRI and neuronal signals (Schölvinck et al. 2010).

Given the putative linkage between M/EEG and fMRI signals, the goal of our current study was to investigate the neural substrates of hemineglect using source-space EEG, which reflects genuine neuroelectric activity uncontaminated by vascular dynamics. Specifically, we performed i) functional connectivity comparisons between hemineglect patients and healthy controls, and ii) regression analyses to identify whole-brain connectivity patterns associated with visuospatial deficits (i.e., perceptual omissions during search tasks) in the left and right hemifields, respectively.

## Methods and Materials

### Study Participants

Stroke patients participated after giving their written informed consent. The study was approved by Geneva State Ethics Committee and accorded with the Helsinki declaration. Patients were admitted after a first right cerebral stroke and consecutively recruited from a primary clinical center. We excluded patients with bilateral lesions, previous neurological or psychiatric disorders, impairment in primary visual perception (except partial visual field defect), psychiatric disorders, severe motor difficulties in the right upper limb, pusher syndrome (i.e., contralateral trunk deviation with active resistance to any attempt of external correction), or current psychotropic treatment. In total, we recruited 15 right-hemisphere-lesioned patients (mean age: 63, SD: 8, 2 women, 13 men) who fulfilled these criteria and were prospectively admitted to two primary clinical centers at the Clinique de Réadaptation of SUVA in Sion (/www.crr-suva.ch) and the Foundation Valais de Coeur of Sion and Sierre (www.valaisdecoeur.ch).

Spatial neglect was assessed using a standard clinical battery similar to other research in our group (Saj et al. 2013; Vaessen et al. 2016), and diagnosed when patients demonstrated abnormal performance in the following tests: line bisection (cut-off: rightward deviation > 11%) (Schenkenberg, Bradford, and Ajax 1980a) and target cancellation test (cut-off: left - right omissions ≥ 4 out of 15 omissions) (Gauthier, Dehaut, and Joanette 1989). All stroke patients (n=15) selected for our sample demonstrated some signs of hemineglect according to at least one of these tests (86% of patients had cancellation deficits, 79% had line bisection deficits), as commonly observed after right hemisphere damage (Halligan et al. 1991; Verdon, Schwartz, Lovblad, et al. 2010) but neglect severity varied substantially between patients. Time since stroke onset was 7.5 months on average (SD 5.3 months) and hemianopia or quadranopia was present in 4 out of 15 patients (26%).

All patients underwent structural MRI scans to delineate the location and extent of brain damage. Individual stroke lesions were manually delineated and the group average lesion map was created using the MRICron toolbox (http://people.cas.sc.edu/rorden/mricron). The maximal lesion overlap affected posterior parts of the lateral prefrontal cortex, the anterior and middle temporal lobe, as we all as the deep paraventricular white-matter in the right hemisphere (see Supplementary Fig S1).

As we sought to explore electrophysiological differences between stroke patients and the healthy population, we also collected data from a control group of 27 matched healthy adults (mean age: 56, SD: 7, males: 23, females: 4), randomly sampled from the Human Brain Institute (HBI) normative database (http://www.hbimed.com/) (Grin-Yatsenko et al. 2009). Importantly, the HBI data were collected using the same EEG amplifier type (Mitsar-201) for recordings of healthy subjects as for the stroke patients.

### Clinical battery for visuospatial neglect

A series of standard paper-and-pencil tasks were presented to each patient at their first visit, in order to quantify patients’ baseline visuospatial biases. Neglect severity was measured with the Schenkenberg line bisection task (Schenkenberg, Bradford, and Ajax 1980b) (18 horizontal lines,10-20cm) and a variant of the bell cancellation test (Gauthier et al. 1989) (35 animal targets among distractor objects) (Gassama et al. 2011). In the latter Animal Cancellation Test, the search sheet was divided into seven virtual columns, each containing 5 targets.

### EEG recording and processing

A multichannel EEG cap was used to measure whole-scalp activity in each baseline recording. This consisted of resting state measurement of 3-minutes under eyes open conditions. All EEG recordings were performed during the clinical workup of patients. Using a similar clinical EEG setup as in preceding work (Tokariev et al. 2018), scalp voltages were recorded with a 19 Ag/AgCl electrode cap (Electro-cap International, Inc. www.electro-cap.com) according to the 10-20 international system. The ground electrode was placed on the scalp, at a site equidistant between Fpz and Fz. Electrical signals were amplified with the Mitsar 21-channel EEG system (Mitsar-201, CE0537, Mitsar, Ltd. http://www.mitsar-medical.com) and all electrode impedances were kept under 5 kΩ. For online recording, electrodes were referenced to linked earlobes, and then the common average reference was calculated off-line before further analysis. EEG data was recorded at 250 Hz and then filtered with a 0.5–40 Hz bandpass filter off-line.

All EEG data were imported into the Matlab toolbox EEGLAB v12 (http://sccn.ucsd.edu/eeglab/) for offline processing. We used Infomax ICA decomposition to remove usual eye movement such as saccades or blinking (Jung et al. 2000). Recordings were further cleaned with an automated z-score based method, using the FASTER plugin (Nolan, Whelan, and Reilly 2010), rejecting 1-second epochs that deviated from the mean by more than 1.5 standard deviations.

### Source-space measures of EEG activity

Artifact-free EEG data were processed in Matlab with the Brainstorm Toolbox (http://neuroimage.usc.edu/brainstorm/). In line with previous approaches using a similar EEG setup in clinical populations (Tokariev et al. 2018), we first computed a head model of the cortex surface for each EEG recording using overlapping spheres (OpenMEEG) and then estimated unconstrained cortical sources using the minimum-norm sLORETA algorithm implemented in Brainstorm. To normalize sources across participants, we projected (warped) the sources from each participant onto the MNI/Colin27 template brain (Collins et al., 1998). The 15,000 voxel source-space was then divided into 68 cortical regions-of-interest (ROIs) according to the Desikan–Killiany neuroanatomical atlas (Desikan et al., 2006). Temporal source-activities across all the voxels in each ROI were then averaged and band-pass filtered in the following 6 frequency bands: delta 1-4 Hz, theta 4-8 Hz, alpha 8-12 Hz, beta 13-30 Hz, gamma 30-45 Hz. For every subject, each frequency band was quantified in Brainstorm to examine differences in spectral power and functional connectivity between the patient and control groups.

#### Spectral power

Band-limited EEG power was estimated with a standard FFT approach using Welch’s method (Matlab pwelch() function) and a Hanning windowing function (4 second epoch, 50% overlap). Relative spectral power (i.e., % power) was calculated as the ratio of the mean power in a specific EEG band and the broadband power (1-45 Hz). Multiple comparison correction was performed using the false discovery rate (FDR) option in the Brainstorm toolbox.

#### Functional connectivity

A single time-course was constructed for each ROI, which was then defined as a *node* in a network graph. Connectivity between nodes was subsequently estimated using the amplitude envelope correlation (AEC), yielding a 68 x 68 node adjacency matrix. The first step in estimating the AEC is to compensate for spatial leakage confounds using a bi-directional orthogonalisation procedure (Hipp et al. 2012) to remove all shared signal at zero lag between filtered EEG signals. After this, we computed the instantaneous amplitude (i.e. envelope) across time for each frequency band by using the absolute value of the Hilbert transform. Finally, the linear correlation between the amplitude time-series of each node pair was calculated using the Pearson coefficient (Brookes et al. 2011). The Fisher r-to-z transformation was applied to the resulting correlation maps.

### Statistical analyses

Source-space (voxel-wise) statistical comparisons of band-limited spectral power were performed using the Brainstorm Toolbox via independent two-tailed t-tests with a p < 0.05 threshold. Separately, we used the GraphVar Toolbox for statistical comparisons of network connectivity using inter-group T-tests with a p < 0.05 threshold. Individual neglect severity was measured by performance on the cancellation test and subsequently used in a between-subject regression analysis (with a p < 0.05 threshold). The left/right visuospatial bias in attention was quantified by the number of omitted targets (i.e. errors) within left/right hemifields, respectively, on the cancellation test.

For all tests, and to statistically correct for multiple-comparisons in functional connectivity measures, we used the Network-Based Statistic (NBS) (Zalesky, Fornito, and Bullmore 2010) based on MATLAB code from the Brain Connectivity Toolbox (brain-connectivity-toolbox.net).We performed n=5000 permutations to generate a null network model based on a random shuffle of all connections. Basically, the NBS is used to control the family-wise error rate when the null hypothesis is tested independently at each of the N(N-1)/2 edges comprising the connectivity matrix. The NBS may provide greater statistical power than conventional procedures such as the false discovery rate (FDR), when the set of edges at which the null hypothesis is rejected constitutes large component(s).

## Results

### Group functional connectivity in patients compared to healthy elderly controls

We defined the EEG source-space functional network connectivity (FNC) graphs for both the stroke patients (n = 15) and the age-matched healthy controls (n = 27), and then contrasted the graphs from each group in order to test for changes in connectivity patterns across the different frequency bands. As illustrated in Fig 1, this revealed three main patterns in the patients relative to controls: i) the delta-band showed a *mixed pattern* of both increased and decreased FNC; ii) the theta-, alpha-, and beta-bands all displayed *decreased* FNC, but with distinctive distributions; and iii) the gamma-band demonstrated selective and localized *increases in* FNC.

**Fig 1.**
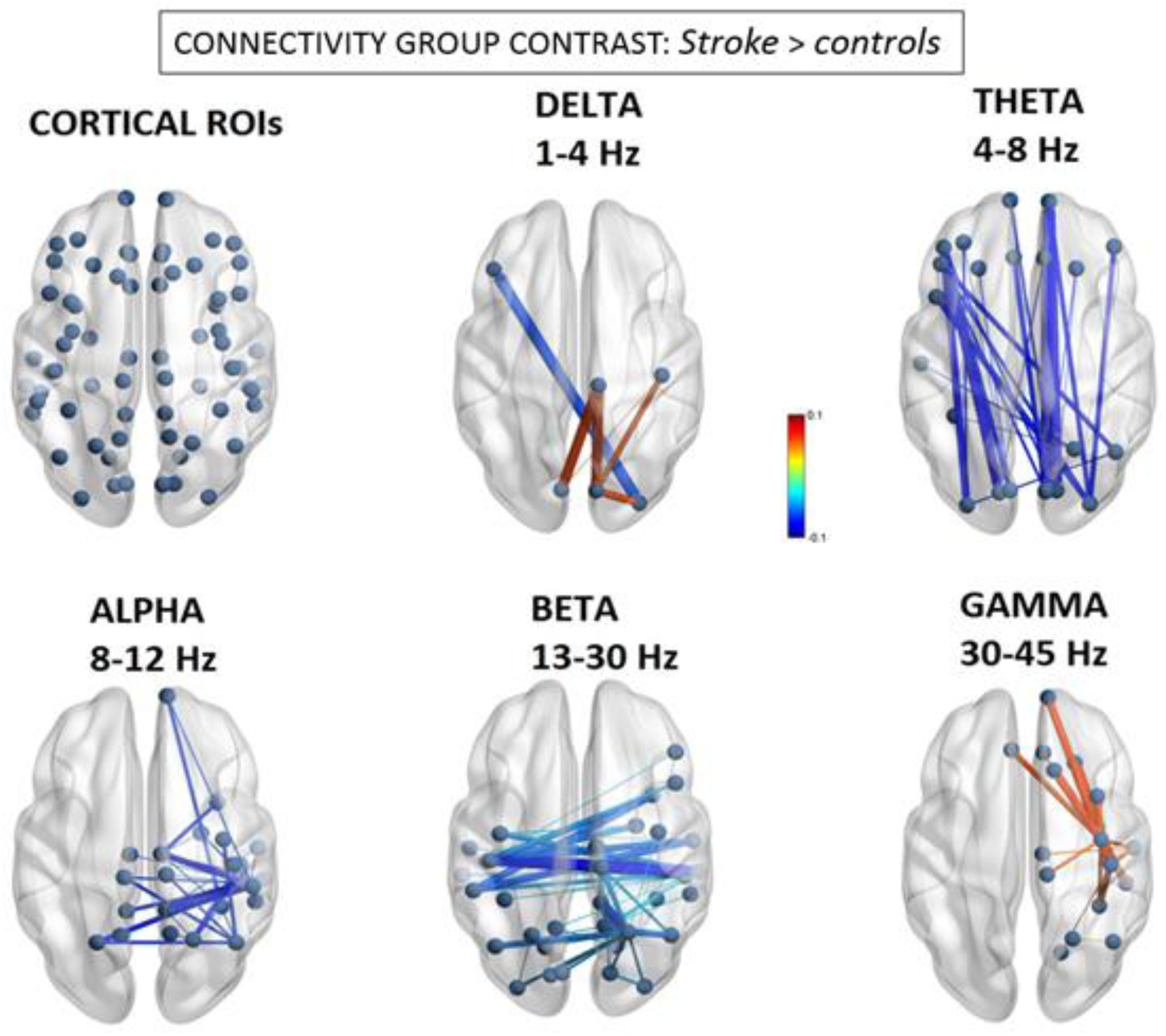
Changes in cortical source functional connectivity. Significant differences of the magnitude of amplitude envelope correlations (AEC) between hemineglect patients and controls. *Red colour* indicates greater connectivity for patients vs controls, while *blue colour* indicates greater connectivity for controls vs patients (p < 0.05 Network Based Statistic (NBS) corrected).

More specifically, using a Network Based Statistic (NBS) correction threshold of p < 0.05, we found the delta-band demonstrated significant but limited changes with mainly intrahemispheric hyperconnectivity, predominating between the right paracentral lobule (BA4) and occipital areas (BA18) (d = 0.063, t = 3.28; d = −0.063, t = 3.58, for sources in right and left cuneus, respectively), but also inter-hemispheric hypoconnectivity between the right lateral occipital cortex (BA18) and left inferior prefrontal cortex / pars triangularis (BA45) nodes (d = −0.064, t = −3.36).

For the theta-band, we observed mainly long-range, intra-hemispheric hypoconnectivity changes along the anterior-posterior axis that, remarkably, were present on both sides. These decreases were the largest between the left pericalcarine (BA17) and left inferior prefrontal gyrus / pars orbitalis (BA47) nodes (d = −0.079, t = −4.51), as well as between the right pericalcarine (BA17) and right frontal pole (BA11) nodes (d = −0.081, t = −4.15).

In sharp contrast, the beta-band showed an opposite pattern of hypoconnectivity mainly affecting interhemispheric connections, particularly those centered on posterior RH areas. These interhemispheric changes affected connectivity between homologous areas of the superior parietal and temporal cortices. Here, the greatest reductions in connectivity were found within the right hemisphere, between the right superior parietal lobule (BA7) and left posterior cingulate gyrus (d = −0.051, t = −5.59), as well as between the right middle temporal (BA20) and right precuneus (BA3) nodes (d = −0.041, t = −4.93).

Alpha-band hypoconnectivity was characterized by a more focal cluster of reduced connections emerging from the posterior right hemisphere. The most salient inter-hemispheric dysconnectivities were observed between right posterior temporal (BA20) and left superior parietal (BA7) nodes (d = −0.130, t = −4.65), as well as between right inferior parietal (BA39) and left superior parietal (BA7) nodes (d = −0.126, t = −5.32).

Finally, the gamma-band displayed a local hyperconnectivity pattern predominating mainly within the anterior right hemisphere. These difference were maximal between the right precentral (BA6) and right superior frontal (BA32) gyri (d = 0.065, t = 2.50), between the right precentral (BA6) and right inferior temporal (BA20) gyri (d = 0.064, t = 2.30), as well as between the right precentral (BA6) and right frontal pole (BA10) (d = 0.062, t = 2.35).

### Relationships between network connectivity and visuospatial bias in hemineglect patients

Although group-level analyses indicated anomalous FNC between patients and control subjects, they do not disambiguate which network connections, if any, may be related to the emergence of visuospatial deficits. All patients presented signs of hemispatial neglect, as commonly observed in right brain-damaged patients (Verdon, Schwartz, Lovblad, et al. 2010), but to a varying degree. The FNC changes described above could more generally reflect the effect of posterior stroke damage. Hence, in order to disentangle neurobehaviorally specific from non-specific connections in stroke-associated hemineglect, we performed regression analyses directly testing for any relation between individual patients’ FNC matrices and the severity of their neglect symptoms, as indexed by the number of left/right omission errors in the visual cancellation test.

Using a Network Based Statistic (NBS) correction threshold of p < 0.05, we found significant brain-behavior relationships that distinctively affected the alpha, beta, and gamma bands. Changes in the delta and theta bands did not predict the severity of visuospatial neglect.

As seen in Fig 2, the alpha-band demonstrated a remarkable contrast between omissions in left and right space, implicating near mirror-image interhemispheric connections. Consistent with the group-level hyponnectivity described above, the number of leftward misses was associated with selective *decreases* in connectivity between extrastriate visual regions in the right inferior temporal gyrus (BA20) with the left precuneus (BA3) (beta = −0.99) and left pericalcarine region (BA17) (beta = −0.98). Conversely, however, the number of misses within the right hemifield (typically reflecting more extensive spatial neglect) was significantly predicted by *increases* in alpha-band connectivity within a homologous set of visual and parietal nodes which included the left inferior temporal cortex (BA20) and the left precuneus (BA3). Importantly, a separate regression with the *total number of left and right misses* (see Supplementary Fig S2) did not reveal any shared connections with either left or right omissions, but limited reductions in connectivity between left medial parietal and right inferior frontal areas, reinforcing the idea that the changes in alpha-band may be specifically related to spatially directed attentional processing.

**Fig 2.**
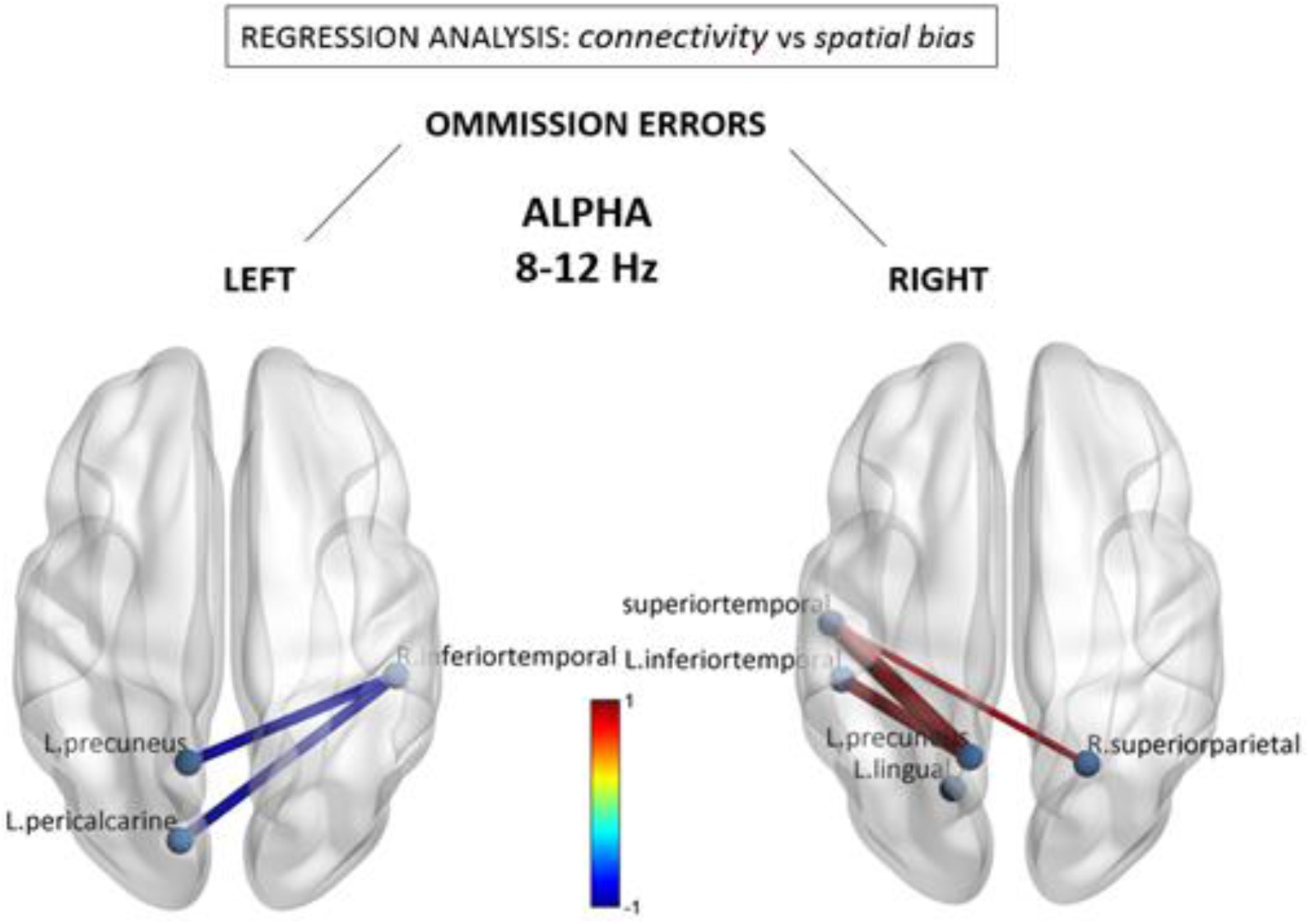
Alpha (8-12 Hz) connectivity as a function of neglect symptoms. Depicted connections correspond to changes in individual pairwise FNC correlating with the number of omissions in the left and right hemifields, respectively, during the cancellation task. Red/blue values indicate statistically significant beta coefficients (p < 0.05 Network Based Statistic (NBS) corrected). Further conjunction analysis (see text) revealed that reduced connections associated with left omissions (in left panel) were those showing the most significant overlap with general stroke-related decreases at the group level.

Fig 3 shows results from similar regression analyses for the beta-band, demonstrating an association of leftward omissions with a different pattern of reduced functional connections between the right posterior superior temporal region (BA22) and anterior brain regions in the left middle frontal gyrus (BA6) (beta = −0.67) and left inferior frontal gyrus / pars orbitalis (BA47) (beta = −0.70). Reductions were also seen within the anterior left hemisphere. No significant associations were observed for the number of rightward errors, nor were there any systematic functional connectivity changes that regressed with the total number of left and right omissions (see Supplementary Fig S3).

**Fig 3.**
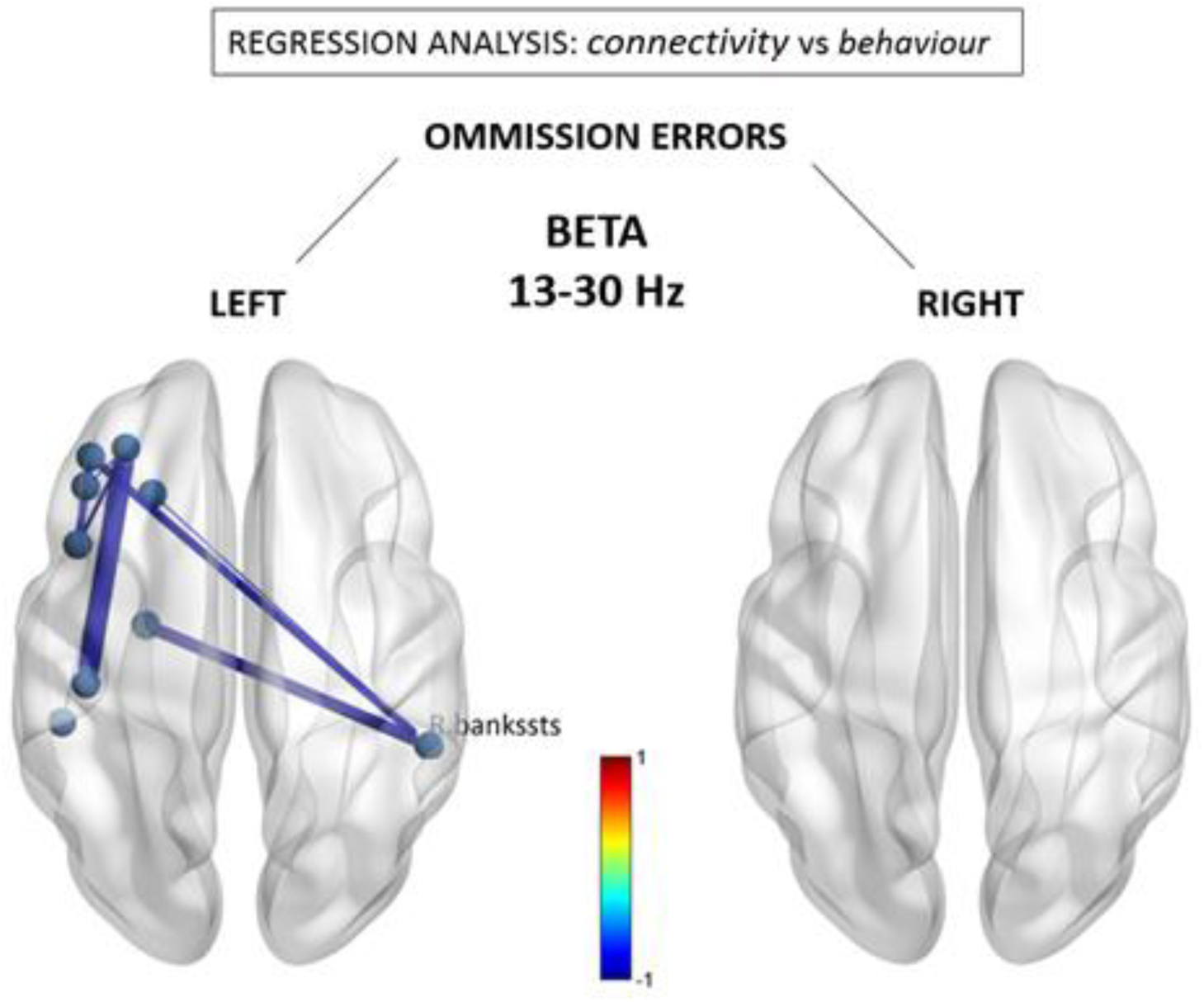
Beta (13-30 Hz) connectivity as a function of neglect symptoms. Depicted connections correspond to changes in individual pairwise FNC correlating with the number of omissions in the left and right hemifields, respectively, during the cancellation task. Red/blue values indicate statistically significant beta coefficients (p < 0.05 Network Based Statistic (NBS) corrected)

Lastly, the relationship between leftward omissions and FNC within the gamma-band revealed a node also implicated within the beta-band, namely the right posterior-superior temporal region (BA22) that acted as a hub for reduced connections with mirror areas in left temporal regions. Also similar to the beta-band, additional reductions were seen within the anterior left hemisphere. However, surprisingly, these negative changes contrasted with the overall increases in connectivity observed for the gamma band at the group-level (see Fig 1). There was no correlation between connectivity patterns in gamma-band and omissions in right space.

### Conjunction analysis between group functional connectivity abnormalities and correlates of visuospatial errors

Finally, we asked whether abnormal functional connections seen at the group level are consistent with those that inter-individually predict stronger leftward visuospatial bias. In other words, is there evidence for a selective disruption of FNC pathways associated with the syndrome of hemineglect? Hence we performed a conjunction of tests (i.e. global null hypothesis) to identify the overlap - if any - between the statistically-significant (p < 0.05 corrected) group-level FNC difference maps (i.e. Fig 1) and the counterpart visuospatial regression maps (i.e. Figs 2 and 3), across the statistically-significant alpha, beta, and gamma bands.

Interestingly, we identified a single overlap with a node in the *right inferior temporal gyrus (BA20) within the alpha band*, which was inter-hemispherically disconnected with the left precuneus (BA3) and left pericalcarine region (BA17); identical to the connections illustrated in the left panel of Fig. 2. This suggests that among all connections globally disrupted by right hemispheric stroke, these may be the most reliable marker of left hemineglect symptoms.

### Group spectral power in patients compared to healthy elderly controls

#### Absolute power

In addition to inter-regional FNC measures, regional source spectral-power (i.e. current source density) differences were estimated between stroke patients and controls using independent t-tests across the five EEG bands (FDR-corrected). Absolute power was consistently *more elevated* in the right (damaged) hemisphere of patients relative to controls for all EEG bands (delta, theta, beta, gamma) *except for alpha,* where no significant differences were detected (p<0.05 corrected). As seen in Supplementary Fig S4, absolute delta power was stronger within posterior parietal areas (t = 3.1, p < 0.05 corrected), theta power within widespread regions with a maximum over supramarginal regions (t = 3.8, p < 0.05 corrected), while beta was higher mostly over frontal (t = 2.7, p < 0.05 corrected) and gamma mostly over precentral motor regions (t = 3.8, p < 0.05 corrected). No significant associations were detected between absolute EEG power changes and visuospatial performance (p < 0.05 corrected).

### Relative power

We also tested for differences in relative (%) power, often used to normalize spectra under a constant value of broadband (1-40 Hz) power, and reflecting the degree of spectral slowing (i.e. greater relative power in lower frequencies) or spectral acceleration (i.e. greater relative power in higher frequencies). As evidenced by Supplementary Fig S5, neglect was associated with relative spectral slowing of right posterior cortical regions, with relative delta power being more elevated in patients mainly within the inferior parietal / posterior temporal junction (t = 3.6, p < 0.05 corrected) while relative theta power was higher within the superior parietal cortex (t = 3.4, p < 0.05 corrected). Conversely, relative alpha power was maximally reduced within the inferior parietal / posterior temporal junction (t = −3.6, p < 0.05 corrected), in conjunction with relative beta (t = −4.4, p < 0.05 corrected) and relative gamma (t = −3.2, p < 0.05 FDR corrected) reductions within the superior parietal lobe. No significant associations were detected between relative EEG power changes and visuospatial performance (p < 0.05 corrected).

### Relating connectivity and oscillatory power changes with lesion topology

Given that changes in FC and source-power were topographically-specific, a reasonable question to ask is whether connectivity differences could be explained firstly by changes in signal power? According to this hypothesis, reduced connectivity would indirectly reflect a degraded signal-to-noise ratio caused by a loss of source EEG power. The latter could result in particular from neuronal loss at the site of the stroke lesion. However, additional analyses allowed us to rule out this hypothesis, given the rather limited overlap between anatomical regions with maximal lesion extent and those with strongest disruptions of connectivity. As shown in Supplementary Fig S6, the location of largest lesion overlap occurred in fronto-insular cortex (which was common to no more than 10 patients), encompassing regions such as BA44, BA45, and BA6. This co-localised with 5 RH nodes/parcels of the Desikan-Kiliani atlas used for FNC calculations: *entorhinal, insula, temporal pole, superior temporal*, and *precentral* regions (see supplementary Fig 4 for their exact locations in MNI space). Importantly, removal of these particular nodes (or their connections) from our analysis did not alter our principal findings, which highlight alpha/beta dysconnectivity with more posterior temporo-parietal foci (see supplementary Fig 4 for anatomical verification).

**Fig 4.**
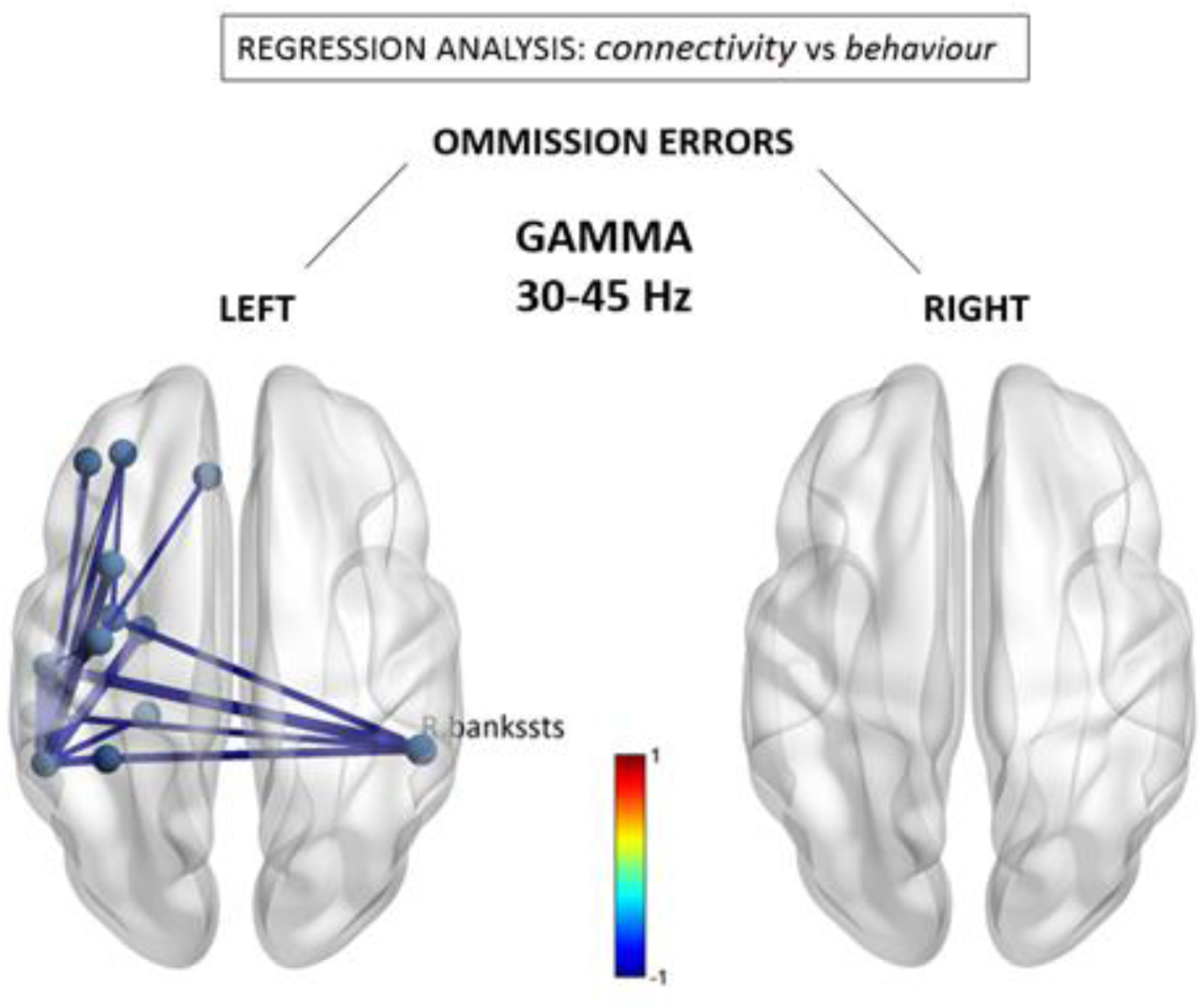
Gamma (30-40 Hz) connectivity as a function of neglect symptoms. Depicted connections correspond to changes in individual pairwise FNC correlating with the number of omissions in the left and right hemifields, respectively, during the cancellation task. Red/blue values indicate statistically significant beta coefficients (p < Network Based Statistic (NBS) corrected)

On the other hand, although regions with significant *relative* power differences (indicative of relative frequency shifts in EEG power) did partly overlap with temporo-parietal nodes presenting abnormal FC power in alpha/beta bands (see supplementary Fig 1), this was clearly not the case for *absolute* source power (see supplementary Fig 2), which provides a more direct measure of electrical signal strength. Notably, each narrowband EEG frequency range (from delta to gamma) exhibited a unique topographical fingerprint (e.g., see Fig. 2), arguing against a simple generative mechanism of reduced EEG signal-to-noise ratio due to neuronal loss that would presumably lead to a broadband attenuation of electrical activity across the full EEG frequency spectrum.

Instead, and given the partial FC overlap with nodal reductions in relative alpha/beta power, we suggest that this pair of EEG rhythms may act as carrier-waves for attentional information (Wróbel 2000) compared to other frequency channels (e.g. delta, theta, gamma), and that it is their *relative* degradation compared to other frequencies that may be an index of impaired network communication (Akam and Kullmann 2012).

## Discussion

Overall, our EEG results converge with previous M/EEG studies reporting electrocortical abnormalities in stroke patients with hemineglect (Colson et al. 2001; Demeurisse, Hublet, and Paternot 1998; Ros et al. 2017; Yordanova et al. 2016) and add novel neurophysiological evidence to support recent findings of neurometabolic (i.e. fMRI) alterations in functional brain networks after stroke (Siegel et al. 2016). Remarkably, we found widespread changes of FNC extending far beyond focal areas of structural brain lesions in RH, including connections with the opposite/intact LH, with very distinctive patterns across different frequency bands. Moreover, a conjunction analysis that sought to overlap i) group-level functional connectivity (FC) differences between patients and controls, together with ii) connections predicting the magnitude of leftward visuospatial bias in a regression analysis, revealed a circumscribed reduction of interhemispheric FC between the right temporo-parietal junction (TPJ) and left precuneal/pericalcarine regions. This is consistent with and supportive of anatomical lesion studies that have long implicated right posterior temporal and inferior parietal areas (in and around the TPJ) as a critical site associated with visuospatial attention deficits (Chelazzi et al. 1998; Heilman and Van Den Abell 1980; Hillis et al. 2005; Karnath, Ferber, and Himmelbach 2001; Mort et al. 2003; Ptak and Valenza 2005; Vallar and Perani 1986), and with the notion of an impaired interhemispheric balance in the pathological spatial biases associated with left hemineglect (Corbetta and Shulman 2011; Kinsbourne 1977; Mesulam 1981). Taken together, our results therefore support the theory that network disturbances characterized by a weakened functional communication between several key cortical regions is a fundamental mechanism underlying spatial neglect (Doricchi et al. 2008; Driver and Vuilleumier 2001; Vuilleumier et al. 2001, 2008), including both interhemispheric and intrahemispheric dysconnectivity.

### The multiplex FNC signature of hemineglect

Compared to age-matched healthy subjects, our source-space analyses of amplitude envelope correlations (AEC) indicated that patients with right hemisphere stroke displayed a multiplex re-organization of neural connectivity that spans the full frequency range captured by EEG, involving delta, theta, alpha, beta and gamma rhythms, and that covered both hemispheres in a frequency-specific manner. Specifically, delta FC was mainly elevated within posterior right hemisphere (RH) regions, while it was reduced between the posterior RH and anterior left hemisphere (LH). These changes are consistent with non-specific effects of brain lesions on focal delta activity (van Dellen et al. 2013), and they did not correlate with neglect symptoms. On the other hand, theta connectivity mainly exhibited a bilateral reduction of long-range intra-hemispheric connections, partially consistent with recent work reporting reductions in fronto-parietal theta and beta coherence in other EEG studies of stroke patients with hemineglect (Fellrath et al. 2016; Yordanova et al. 2017). However, we found that the theta changes did not correlate with neglect severity either, suggesting that the previously reported changes may constitute non-specific sequels of stroke lesions.

In contrast, we found that alpha and beta rhythms both demonstrated important reductions in intra-and inter-hemispheric connectivity of posterior temporal and parietal cortices, i.e., regions traditionally found to be implicated in hemineglect (Thimm, Fink, and Sturm 2008; Umarova et al. 2011; Verdon, Schwartz, Lovblad, et al. 2010). Furthermore, some of these connections were significantly associated with neglect symptoms, in particular those linking right temporo-parietal areas with left posterior parietal regions in the alpha band and with left prefrontal regions in the beta band (see figs 2 and 3). Remarkably, changes in alpha-band connections significantly correlated with the severity of visuospatial biases and implicated symmetrical regions over the extra-striate visual cortex (pericalcarine and lingual), where hypoconnectivity with the left-hemisphere predicted more left visual-field omissions, while the converse (hypoconnectivity with the right-hemisphere) was observed for right visual-field omissions. This topographically-specific dysconnection pattern is compatible with the role of alpha rhythms in contralateral visuospatial attention (Ikkai, Dandekar, and Curtis 2016; Lobier, Palva, and Palva 2017; van Schouwenburg, Zanto, and Gazzaley 2017), as well as previous studies on hemineglect utilizing functional (Ramsey et al. 2016; Sasaki et al. 2013) and structural (Karnath et al. 2011; Vaessen et al. 2016) neuroimaging to delineate the functional neuro-anatomy of visuospatial deficits. Moreover, it is interesting to note that while the alpha band was implicated in the dysconnectivity pattern of right TPJ with sensory visual areas, the beta band was distinctively involved in dysconnectivity with left prefrontal areas which presumably subserve more executive processes of attention (Antzoulatos and Miller 2016).

Importantly, a selective depression of inter-hemispheric connectivity has in itself been shown to be a pathological hallmark of hemineglect in both fMRI and EEG studies (Ramsey et al. 2016; Sasaki et al. 2013), including for the alpha (Sasaki et al. 2013) and beta bands (Guggisberg et al. 2014). Furthermore, increased alpha and/or beta FC has been reported to be a predictive marker of clinical status after stroke or traumatic brain injury, in patients with motor (Dubovik et al. 2012; Guggisberg et al. 2014; Kawano et al. 2017) as well as language (Castellanos et al. 2010; Guggisberg et al. 2014; Nicolo et al. 2015) impairments.

Finally, we also found connectivity changes in the gamma range, including increases within right frontal areas at the whole group level, as well as decreases between right temporo-parietal and left hemisphere regions that correlated with neglect symptoms and partly overlapped with beta changes. These divergent effects make it difficult to associate them with a clear functional role in spatial deficits after stroke. Moreover, gamma-range neuronal activity is usually associated with local interactions between nearby cortical populations rather than with long-distance interactions at network level (Vinck et al. 2013). Given theoretical proposals linking gamma activity to conscious perceptual processes (Melloni et al. 2007) one tentative hypothesis might be that a reduction of synchronous activity in gamma band between right TPJ and left fronto-temporal networks would reflect the loss of access of spatial representations held in the right hemisphere (Karnath et al. 2001) to left hemisphere processes mediating conscious awareness and verbal report, in line with work in split-brain patients demonstrating an intimate link between conscious behavior and language abilities of the left hemisphere(Volz and Gazzaniga 2017). However, this interpretation is speculative and the role of gamma-range connectivity in neglect, if any, remains unsettled and undoubtedly needs further investigation to be clarified.

### Limitations

As already noted above, connectivity measures might potentially be contaminated by degraded signal-to-noise ratio and global losses in source EEG power due to neuronal tissue damage. However, as shown by our supplementary analyses, we consider this interpretation unlikely for several reasons, most notably because connectivity changes included both decreases and increases, they did not correspond to the local peaks of *absolute* source power that constitutes a more direct measure of electrical signal strength, and distinct patterns were observed for each EEG frequency sub-band. Taken together, this argues against a single, global mechanism of reduced EEG signal-to-noise ratio due to neuronal loss. Instead, and given the partial FC overlap with nodal reductions in relative alpha/beta power, we suggest that this pair of EEG rhythms may act as major carrier-waves for attentional information (Wróbel 2000), distinct from other frequency channels (e.g. delta, theta, gamma), and hence it is their *relative* degradation compared to other frequencies that may constitute a functional index of impaired network communication (Akam and Kullmann 2012).

Other potential limitations are reflective of the imaging modality we used, i.e., EEG. Although EEG provides a direct measure of neural activity, it is most sensitive to sources within the cortical-mantle. Hence, our analyses were restricted to cortical network dynamics and did not allow for reliable assessment of subcortical structures which may an important role in the control of attention (Gitelman et al. 1999; Vuilleumier 2013). Another weakness involves the relatively low number of EEG channels that constitute the 10-20 montage, raising potential concerns of excessive source localization error and/or spread, which would lead to spurious activity mixing between cortical ROIs. Firstly, given the clinical nature of our study, we found that the 10-20 montage met an important trade-off between EEG set-up time and patient fatigue during clinical test batteries, which were often flanked by daily rehabilitation sessions. Secondly, our methodology allows for replication and application in standard clinical settings where high-density EEG is not routinely available. Thirdly, a source-localisation simulation study with sLORETA using 1000 cortical sources (i.e. patches) of approx. 1cm^2^ (Song et al. 2015), and an electrode montage based on the 10-20 system, indicated a mean localization accuracy of ∼0.5 cm and source spread of ∼0.7 cm, which remains within the lowest inter-ROI distances of the Desikan-Killiany atlas (∼1.5 cm) used in the present study. Importantly, as the Desikan-Killiany atlas contains ROIs of variable patch sizes, empirical work has not revealed a systematic bias between patch size and sLORETA reconstruction accuracy (Cosandier-Rimélé et al. 2017). Lastly, in line with published work leveraging a similar low-density EEG montage in clinical conditions (Tokariev et al. 2018), our analysis pipeline included EEG signal orthogonalisation prior to FC estimation, which has been shown to minimize any instantaneous (i.e at zero-phase lag) activity that is spuriously shared between ROIs and that may have arisen from (low-resolution) source blurring (Brookes et al. 2011).

## Conclusion

In a nutshell, our results show that the hemineglect syndrome following RH stroke is associated with a widely distributed but anatomically-specific disruption of cortical network connectivity that involves temporo-parietal regions of the right hemisphere and their functional interactions both within and between the two hemispheres. In particular, losses in the connectivity of the posterior RH with left parieto-occipital cortex in alpha channels and with left prefrontal cortex in beta channels appear to be critically involved in impaired control of visuospatial attention towards the contralesional / left hemispace. Better mapping and understanding these neurophysiological markers of hemineglect is an important step towards designing novel tools to assess post-stroke deficits in patients and more effective rehabilitation approaches, notably based on neurofeedback (Ros et al. 2017; Saj et al. 2017) or noninvasive brain stimulation (Lee et al. 2013; Sunwoo et al. 2013).

## Acknowledgements

This study was generously supported by grants from the Leenaards Foundation, as well as the EU Marie-Curie COFUND program BRIDGE (Grant no. 267,171), Geneva University Hospitals (HUG), Société Académique de Genève, and the Swiss National Science Foundation (SNF, Grant no. 320030-166704).

## Supplementary Figures

**Supplementary Figure S1:**
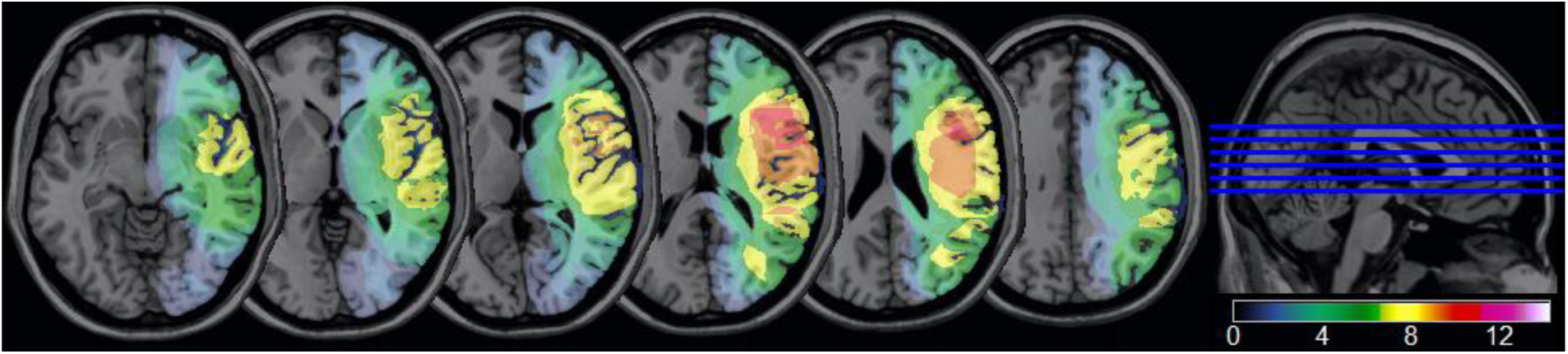
**Average structural MRI lesion maps of the 15 stroke patients** with hemineglect. Colors code for the number of patients with damage to a particular voxel location. Lesions were widely distributed in the right hemisphere, with the strongest overlap in fronto-insular regions (red colour, peak coordinates [x= 42, y= −2, z= 14]), affected in a maximum of 10 patients.

**Supplementary Figure S2:**
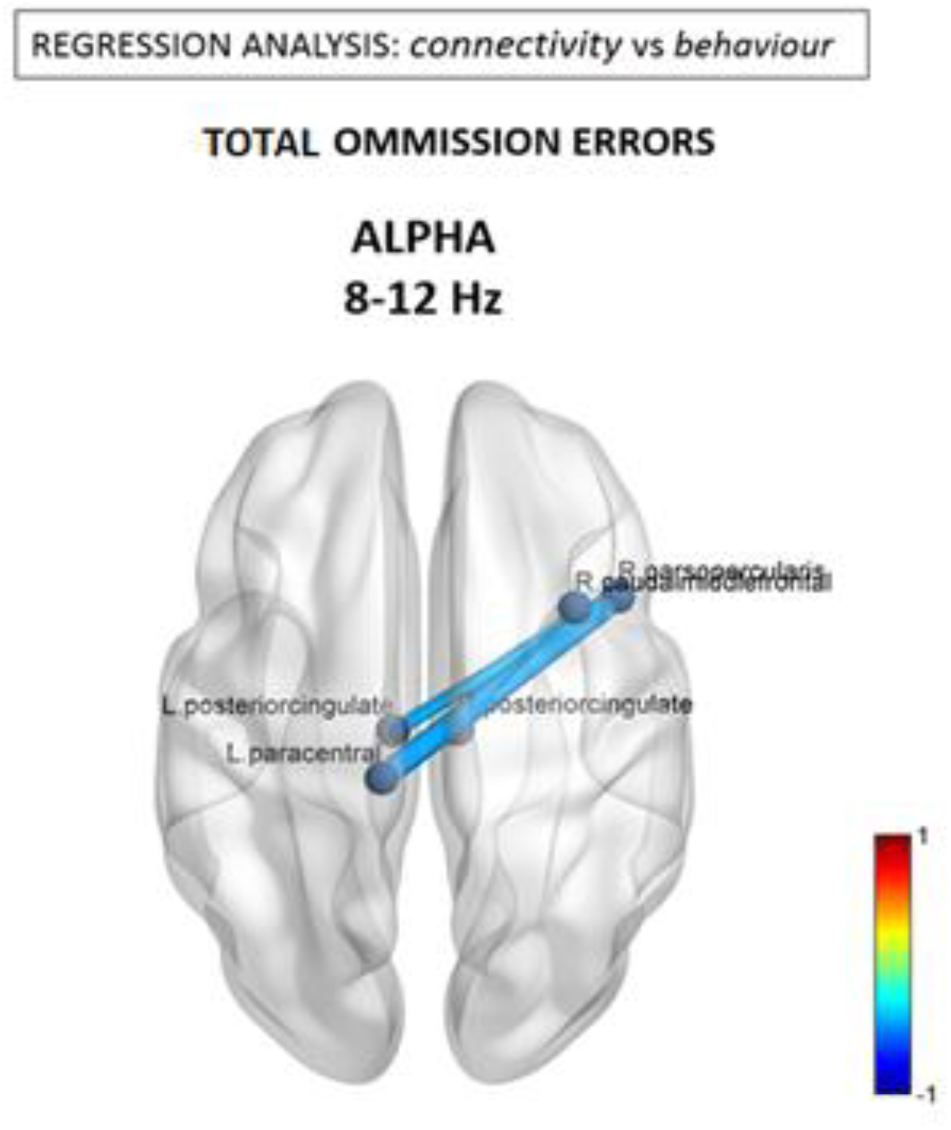
Alpha (8-12 Hz) connectivity as a function of neglect severity. Depicted connections correspond to changes in individual pairwise FNC correlating with the *total* number of left and right omissions during the cancellation task. Red/blue values indicate statistically significant beta coefficients (p < 0.05 Network Based Statistic (NBS) corrected)

**Supplementary Figure S3:**
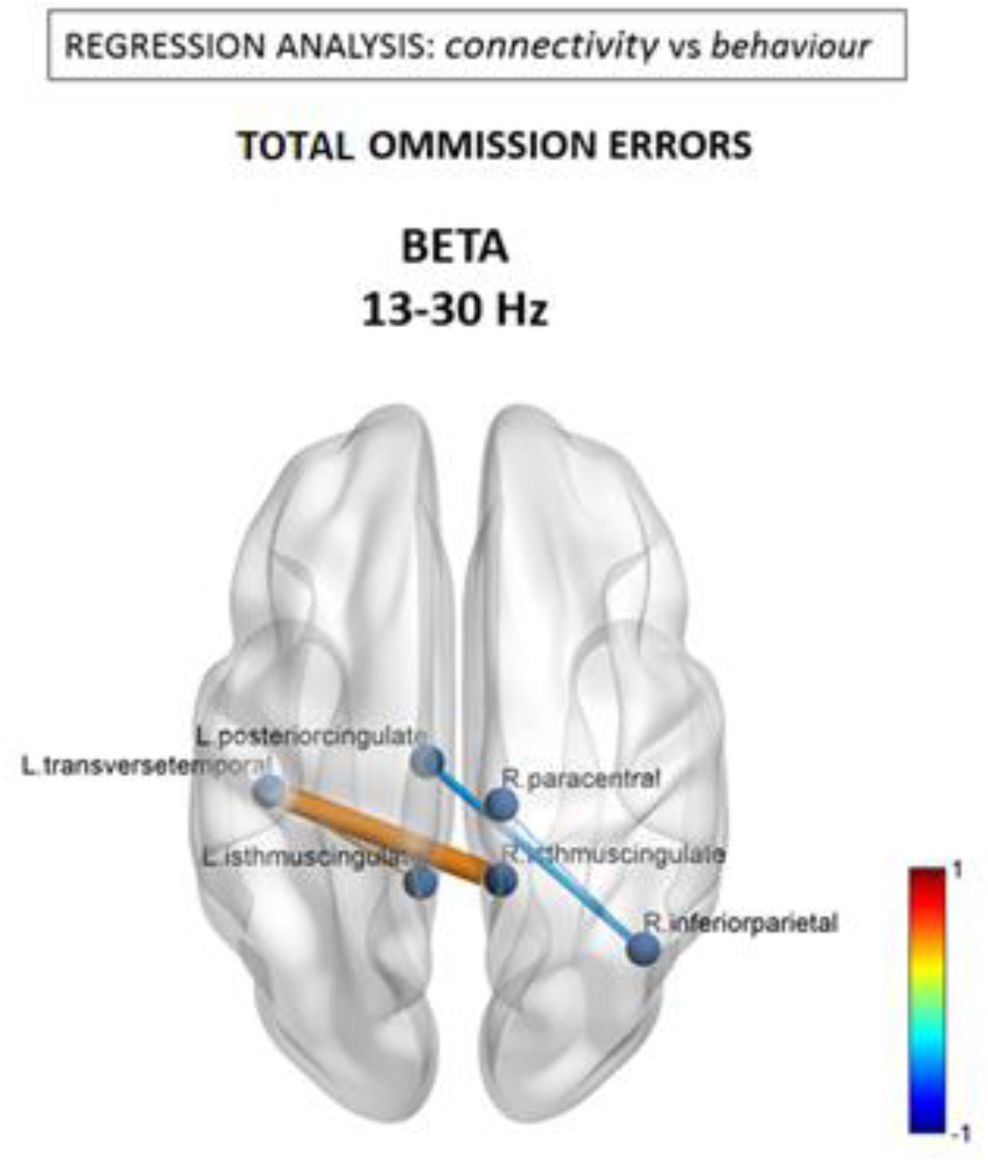
Beta (13-30 Hz) connectivity as a function of neglect severity. Depicted connections correspond to changes in individual pairwise FNC correlating with the *total* number of left and right omissions during the cancellation task. Red/blue values indicate statistically significant beta coefficients (p < 0.05 Network Based Statistic (NBS) corrected)

**Supplementary Figure S4.**
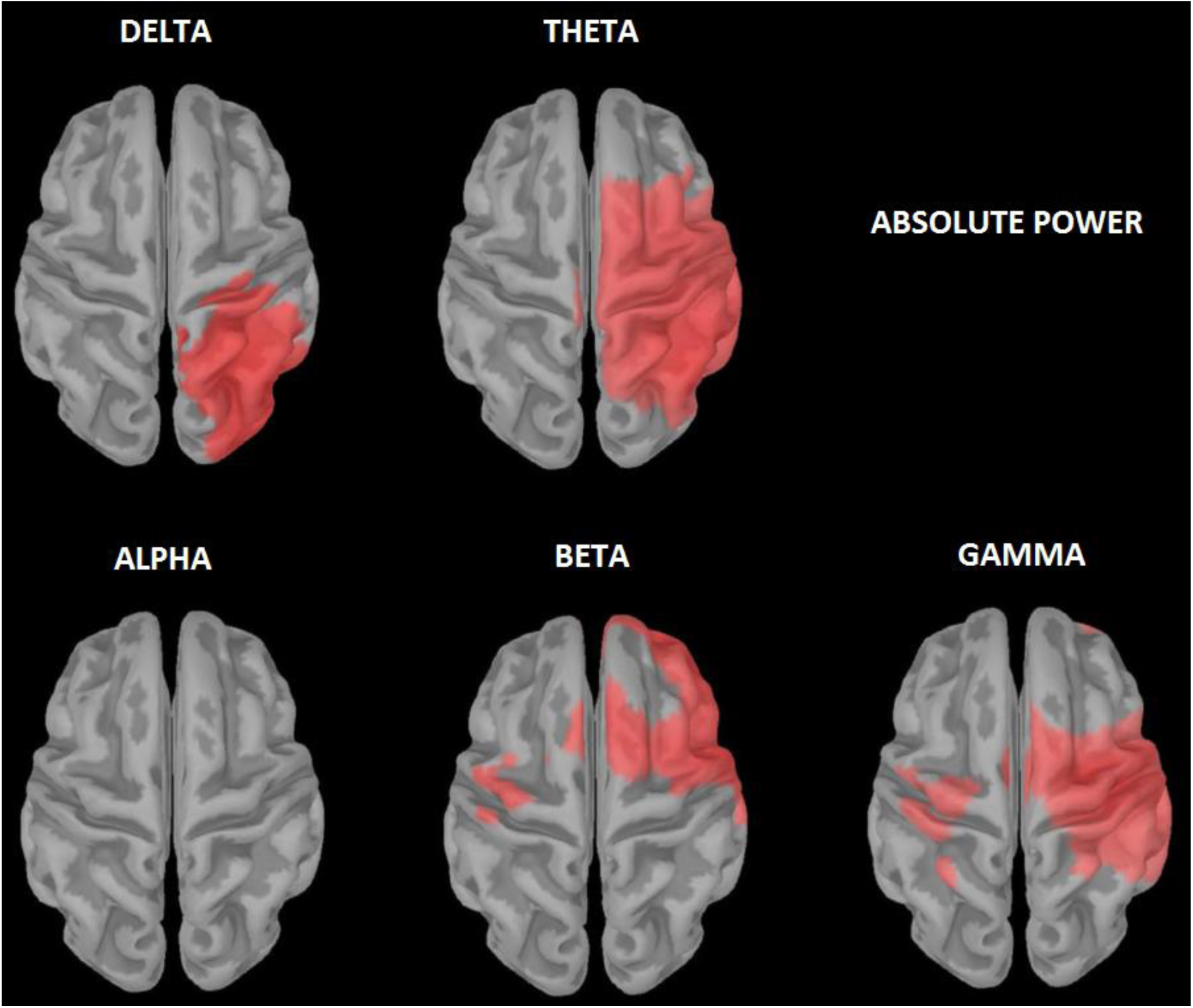
Absolute spectral power. T-values of statistical differences in sLORETA relative source-power between hemineglect patients and controls. *Red colour* indicates greater power for patients, while *blue colour* indicates greater power for controls (p < 0.05 FDR corrected).

**Supplementary Figure S5.**
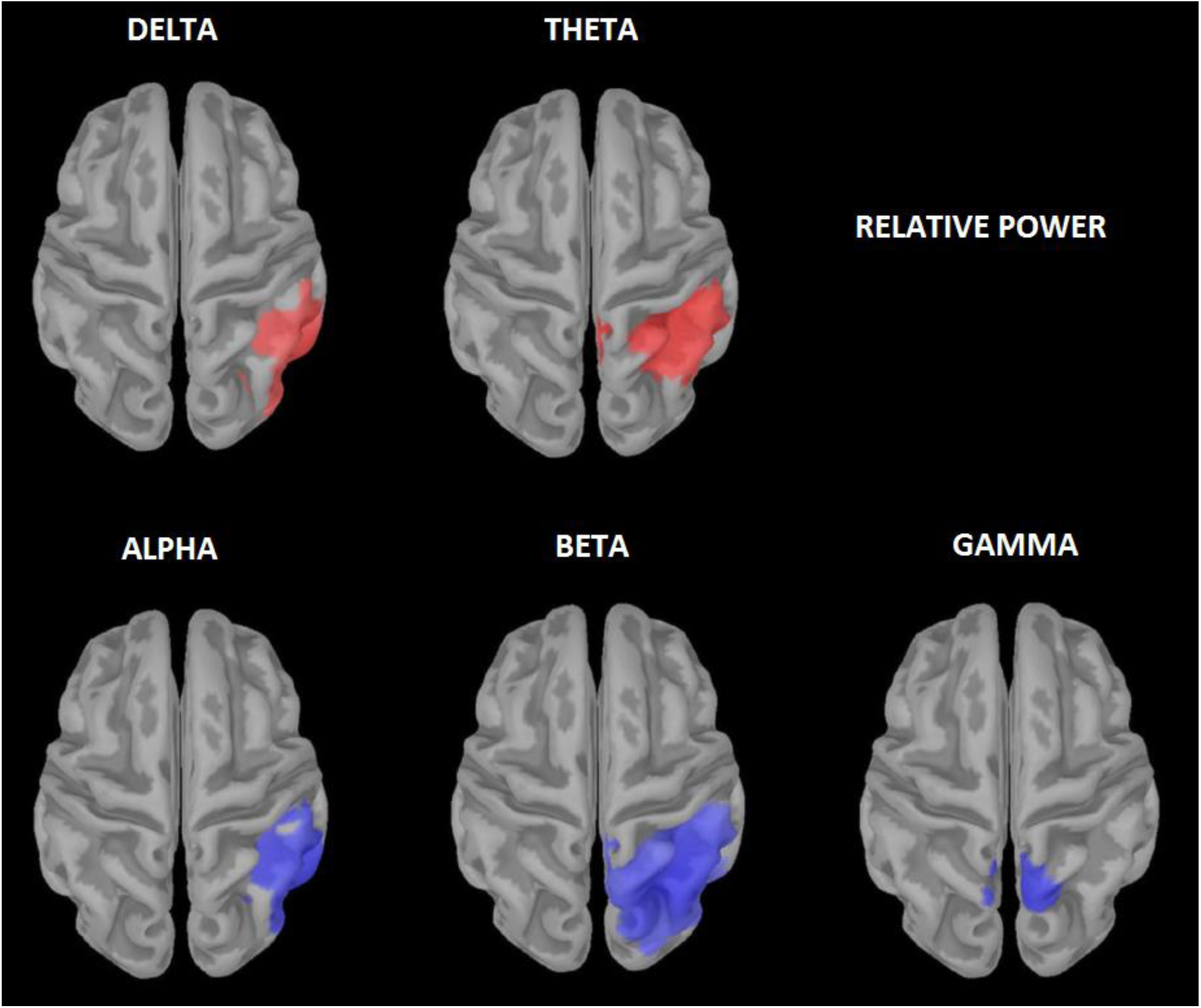
Relative spectral power. T-values of statistical differences in sLORETA relative source-power between hemineglect patients and controls. *Red colour* indicates greater power for patients, while *blue colour* indicates greater power for controls (p < 0.05 FDR corrected).

**Supplementary Figure S6:**
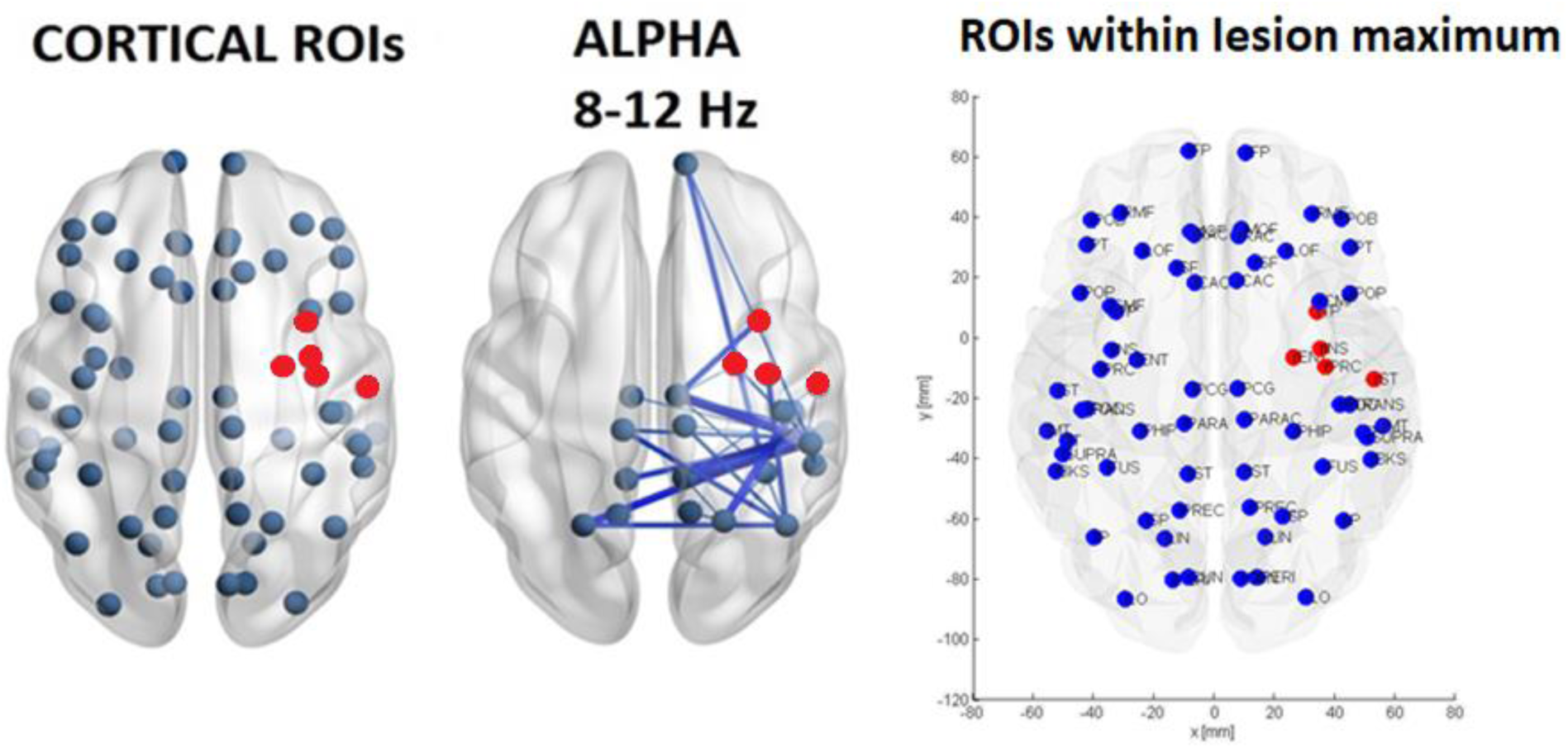
**Network nodes within locus of maximum lesion** with peak coordinates [x= 42, y= −2, z= 14]). Affected nodes are depicted in *red colour* in the 3rd panel with MNI coordinates; 1st and 2^nd^ panels are from Fig 1 for comparison. See also Supplementary Figure 3 for structural MRI image of lesions.

